# Manuscript: Detecting differences in Size Spectra

**DOI:** 10.1101/2023.03.14.532592

**Authors:** Justin Pomeranz, James R. Junker, Vojsava Gjoni, Jeff S. Wesner

## Abstract

1. The distribution of body size in communities is remarkably consistent across habitats and taxa and can be represented by size spectra, which are described by a power law. The focus of size spectra analysis is to estimate the exponent (*λ*) of the power law.
2. Many methods have been proposed for estimating *λ* most of which involve binning the data, summing abundance within bins, and then fitting a ordinary least squares (OLS) regression in log-log space. However, recent work has shown that binning procedures may return biased estimates of size spectra exponents compared to procedures that directly estimate *λ* using maximum likelihood estimation (MLE). Despite this variability in estimates, it is unclear if the relative change across environmental gradients is consistent across methodologies. Here, we used simulation to compare the ability of two binning methods (NAS, ELBn) and MLE to 1) recapture known values of *λ*, and 2) recapture parameters in a linear regression measuring the change in *λ* across a hypothetical environmental gradient. We also compared the methods using two previously published body size datasets across a pollution gradient and a temperature gradient
3. Maximum likelihood methods always performed better than common binning methods, which demonstrated consistent bias depending on the simulated values of *λ*. This bias carried over to the regressions, which were more accurate when *λ* was estimated using MLE compared to the binning procedures. Additionally, the variance in estimates using MLE methods is markedly reduced when compared to binning methods.
4. The uncertainty and variation in estimates when using binning methods is often greater than or equal to the variation previously published in experimental and observational studies, bringing into question the effect size of previously published results. However, while the methods produced different slope estimates from previously published datasets, they were in qualitative agreement on the sign of those slopes (i.e., all negative or all positive). Our results provide further support for the direct estimation of *λ* using MLE (or similar procedures) over the more common methods of binning.

## Introduction

Body size distributions are a fundamental characteristic of communities. In general, size-abundance relationships are expected to scale with the value of -0.75 across all biological communities (Damuth 1981, 1991, Damuth 1998). This is thought to be a consequence of simple size-dependent metabolic constraints on organisms’ energy use predicted by the metabolic theory of ecology (Nee et al. 1991, Brown et al. 2004). The remarkable consistency of these relationships across spatiotemporal scales and ecosystems has led them to be recommended as a “universal” indicator of ecological status (Petchey and Belgrano 2010). Variation in size-abundance relationships have been documented through space (Pomeranz et al. 2022), time (Evans et al. 2022), seasonality (McGarvey and Kirk 2018), and in response to human activities (Jennings and Blanchard 2004). Likewise, variation in size spectra relationships have been used to explain fundamental differences in how communities are organized. For example, external resource subsidies “bend the rules” and allow higher abundances of large body sizes than would be expected (Perkins et al. 2018). However, recent research has shown that these results may be an artifact of how the data were treated. Edwards et al. (2020) reanalyzed a time series of marine fisheries data and found that previously reported changes through time were actually dependent on the methodology used.

Individual size distributions (ISD *sensu* (White et al. 2007), also referred to as abundance size spectra) are one of the size-abundance relationships commonly used. ISDs represent frequency distributions of individual body sizes within a community, regardless of taxonomy. Generally, there is a negative relationship between individual body size (M) on the *x*-axis and abundance (N) on the *y*-axis. Theoretical and empirical data support this relationship being described as a simple power law with exponent *λ* in the form of *N* ∼ *M*^*λ*^, where *λ* = 2 (Sheldon and Kerr 1972, Andersen and Beyer 2006). Commonly, *N* is the count of body sizes grouped into bins, and *λ* is estimated as the slope from OLS regressions in log-log space of *N* and the mid point of the body size bin *M*_*bin*_: log(*N*_*count*_) = *λ* log(*M*_*bin*_). Myriad binning methods have been proposed, including linear and logarithmic bin widths. Likewise, some methods rely on the absolute counts in the bins and others employ normalization techniques such as dividing the count by the bin width (especially common with logarithmic binning). As an alternative to binning methods, *λ* can be estimated directly on un-binned data using maximum likelihood techniques.

Previous work has shown that the estimates of *λ* differ between MLE and size-binned OLS techniques. Size-binned OLS methods are particularly sensitive to decisions made in the binning process. Simulation studies have shown that MLE offers consistently more accurate estimates of *λ* (White et al. 2008, Edwards et al. 2017), and reanalysis of empirical data also indicates that the conclusions are dependent on the methodology used (White et al. 2008, Edwards et al. 2020). However, recent empirical analysis of stream macroinvertebrate communities across the National Ecological Observatory Network (NEON, USA) showed that while the estimates of *λ* varied based on method, the relative change across the environmental gradient was consistent across methods (Pomeranz et al. 2022). While there is a growing consensus that MLE methods offer more reliable estimates of *λ*, and binning methods result in biased estimates, it remains unclear if these biases are consistent and systematic or stochastic, and whether or not the relative change in ISD parameters is consistent across space and time. In other words, if the data within a study are all treated the same, does a relative change of size-binned OLS slope parameters of 0.1 coincide with a relative change of MLE *λ* estimates of 0.1?

We had two primary objectives in this study: 1) to compare how well different methods estimate sitespecific *λ*’s and 2) to compare how different methods of estimating *λ* impact subsequent regressions that use *λ*’s as a response variable. We did this using both simulated and empirical data sets.

## Methods

### Data Simulation

To investigate the performance of commonly used methods, we simulate body size observations from a bounded power law distribution using the inverse method, as described in (Edwards et al. 2017). Let *M* be a random variable of body sizes described by the probability density function:

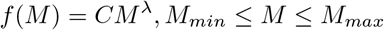

*M*_*min*_ = 0.0026 and *M*_*max*_ = 1.2∗10^3^. These values were based on empirical body sizes of stream benthic communities reported in (Pomeranz et al. 2022). Our results were not dependent on the range of body sizes (Supplemental Information).

### Site-specific *λ* estimates

Using the procedure above, we simulated *n* = 1000 body sizes from five different *λ*’s: (-1.5, -1.75, - 2.00, -2.25, -2.5). The values of *λ* describe how quickly the abundance of large body sizes decline within a community. For example, a value of -1.5 means there would be a relatively high number of large body sizes (shallow) whereas a value of -2.5 means there would be relatively very few large body sizes (steep).

### Estimation of Size spectra parameter *λ*

After simulating data, we used three different methods (described below) to estimate the value of *λ* (maximum likelihood, equal logarithmic bins, and normalized abundance spectra). We repeated the process of data simulation and parameter estimation 1000 times (reps) and plotted the distribution of values obtained for each method.

### Maximum likelihood estimation (MLE)

We estimated the exponent *λ* directly using MLE methods modified from the the sizeSpectra package (Edwards et al. 2017). Throughout the manuscript, these estimates are referred to as MLE.

### Equal Logarithmic Bins: ELBn

In addition, to estimate the OLS slope parameter in log-log space, we used two common binning methods. It is important to note that the log-log regression is estimating *λ* as the regressionslope (*β*_1_), and not *λ* directly. In the interest of clarity, we will refer to all estimates of lambda as *λ* regardless of the method used, and reserve the terms “parameters,” “intercept,” and “slope” *β*_*env*_ to refer to the parameters of linear regressions that use the lambdas as a response variable.

For the first binning method, we created six equal logarithmic bins covering the range of body sizes. This method has been used extensively in previous studies (e.g., see Perkins et al. (2018), Martinez et al. (2016), and Dossena et al. (2012). In the present study, the count in each bin was normalized by dividing by the bin width to account for the unequal bin sizes. The process of normalizing shifts the estimate by -1. In other words, an un-normalized estimate of -0.75 would result in an estimate of -1.75 when normalizing the data (Sprules and Barth 2015, Edwards et al. 2017, Pomeranz et al. 2022). Throughout the manuscript, the normalized equal logarithmic binning method will be referred to as ELBn.

### Log_2_ Bins: NAS

The second binning method was similar to ELBn, but bins of log_2_ width are used. The number of bins is dependent on the range of body sizes present in the data. When working with empirical data with different size ranges this can alter the number of bins per site, and it is recommended to construct the bins based on the global minimum and maximum body size present. However, since the data here were simulated from a known size range, the number of bins for each site is identical. The count in each bin is normalized in the same way as described above for the ELBn approach. This method has been used extensively in the literature (e.g., Jennings et al. (2002, 2004), Sprules and Barth (2015), Pomeranz et al. (2019a), McGarvey and Kirk (2018)), and is referred to as the Normalized Abundance Spectrum (NAS).

For the two binning methods, simple OLS regression were conducted in the form

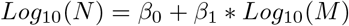

Where *N* is the count in each bin, *M* is the mid-point of each bin, and *β*_1_ is considered to be the *λ* estimate.

### Variation in *λ* Across a Gradient

Previous work has investigated the biases in *λ* estimates when using different methods (White et al. 2007, Edwards et al. 2017, Edwards et al. 2020). However, the focus of the present work is to investigate biases when estimating the relationship parameters across an environmental gradient.

To do this, we first simulated *λ*’s from a linear regression with a known slope *β*_1_ of -0.5 and intercept of -1.5 across a generic predictor variable x with values ranging from -1 to 1:

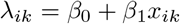

This produced *k* = 1000 values from each value of *x* (−1, -0.5, 0, 0.5, 1), resulting in 5000 total *λ*’s. From each *λ* we simulated 1000 individual body sizes using the procedure described above.

Once the body size data were simulated, we then reversed the procedure. First, we used the three methods (NAS, ELBn, MLE) to estimate *λ* for each of the 5000 body size data sets. Then, we fit a linear regression to those data with the predictor variable *x* and compared the resulting intercept (*β*_0_) and slope (*β*_1_) parameter estimates to the known value. The main results presented here were not dependent on the range of x-values or the number of sites (Supplemental Information).

### Empirical Data

We re-analyzed two data sets of benthic macroinvertebrate communities from stream habitats across two different gradients. In the first, quantitative macroinvertebrate samples were collected from streams across an acid mine drainage (AMD) stress gradient. Details of the sample collection and processing can be found in (Pomeranz et al. 2019a). Briefly, all individuals from each sample were identified to the lowest practical taxonomic unit and body lengths were measured using image processing software from photos taken with a camera mounted to a dissecting microscope. Body mass was estimated using taxon-specific published length weight regressions.

The second data set was from the wadeable stream sites of NEON (National Ecological Observatory Network (NEON) 2022). NEON stream sites are located across a wide temperature gradient in the United States, from Puerto Rico to Alaska. Quantitative macroinvertebrate samples were collected using the most appropriate method based on the local habitat. All individuals were identified and had their body lengths measured, and body mass was estimated using published length-weight regressions. This data has been analyzed previously using size spectra methods as described in Pomeranz et al. (2022). Detailed methods of the sampling collection and data processing methods can be found in the macroinvertebrate data product information documents found on the NEON website https://data.neonscience.org/data-products/DP1.20120.001. Estimates of the slope coefficient (*β*_1_) ± 1 SD, are compared across methods. This allows us to determine whether or not the main results would differ depending on the method used.

### Performance Metrics

We compared performance of each procedure (NAS, ELBn, MLE) by first plotting the distribution of lambda or regression parameter estimates (*λ, β*_0_, *β*_1_) against the known values. We calculated bias for the procedures overall as the median absolute difference (averaged across all simulations) between the known values and the modeled estimates. Finally, for each procedure we estimated the width of the 95% CI’s to compare uncertainty.

### Data Availabity

The empirical data used in this manuscript is already publicly available (Pomeranz et al. (2019b); data dryad DOI https://datadryad.org/stash/dataset/doi:10.5061/dryad.v6g985s, and (National Ecological Observatory Network (NEON) 2022). R Scripts to reproduce the full simulation and analysis are available at: ([https://github.com/Jpomz/detecting-spectra-differences]). (To be archived upon acceptance.)

## Results

### *λ* estimates

There was considerable variation in the *λ* estimate across methodologies (Figure 1). The distribution of estimates from the MLE method was always symmetrical and centered at the known value of *λ* (Figure 1). The distribution of estimates from the binning methods were generally wider and occasionally asymmetrical or bimodal. On average, the 95% confidence intervals produced by the NAS and ELBn methods were ∼2 times wider than those produced by MLE (Table 1), indicating greater consistency of estimates from MLE. Similarly, estimates of *λ* deviated from the true *λ* by an average of 0.04 or 0.05 absolute units for the NAS and ELBn methods, more than four times higher than the deviation (0.01) observed for the MLE (Table 1).

**Figure 1:**
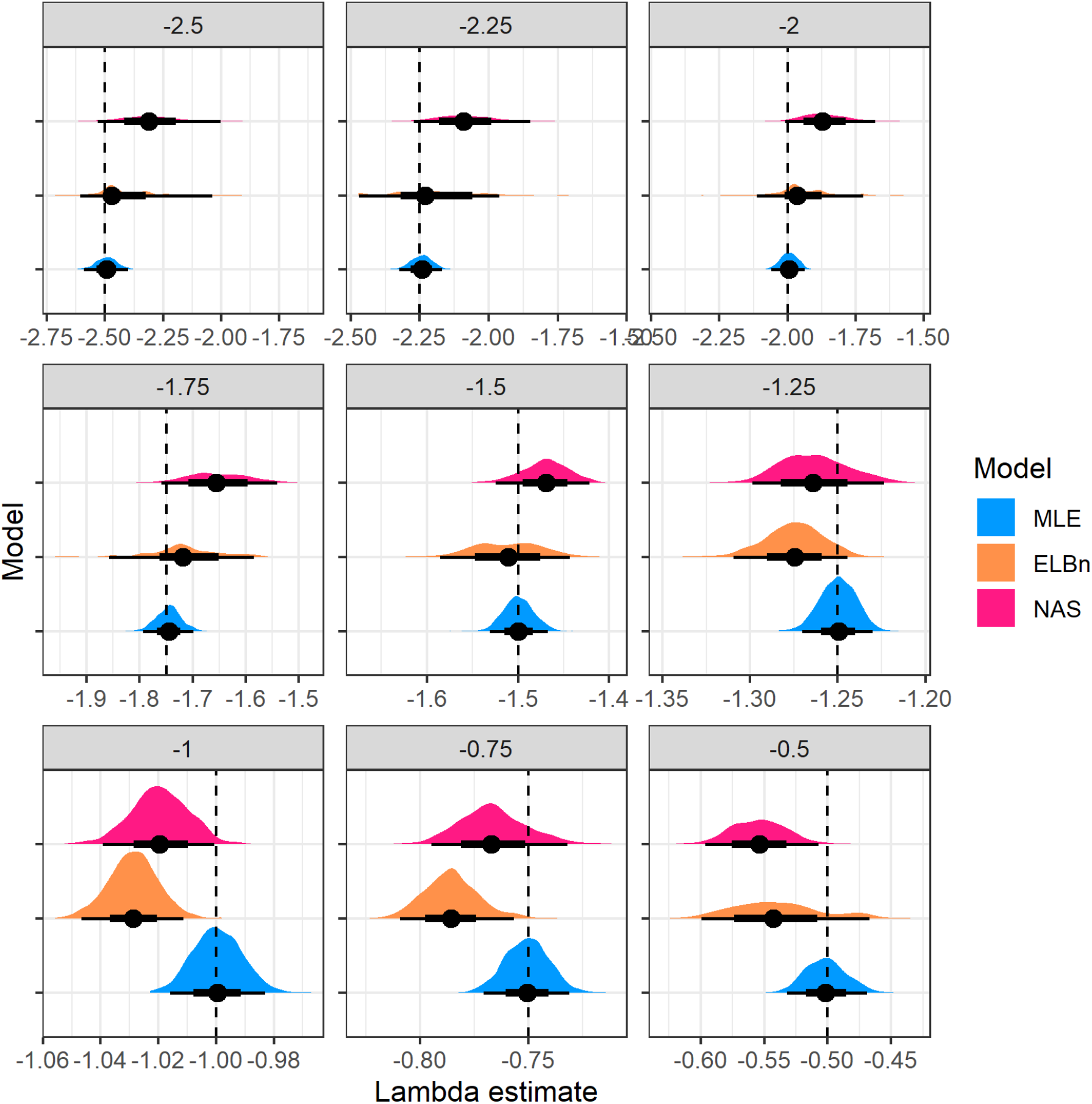
Figure 1. Distribution of Lambda estimates by method (color) from random samples of body sizes from bounded power law distributions with varying exponents (−2.5 to -0.5). The figure is facetted by the known lambda parameter (facet title) and is also shown as the dashed line in each facet. Note that the x-axis varies in each facet.

Furthermore, the binning methods systematically over estimated steep size spectra relationships (*λ* = ∼ - 2.5 *to* - 1.5, Figure 2), and slightly underestimated shallow size spectra relationships (*λ* = ∼ - 0.5). This finding was more pronounced in the NAS method compared with the ELBn method.

**Figure 2:**
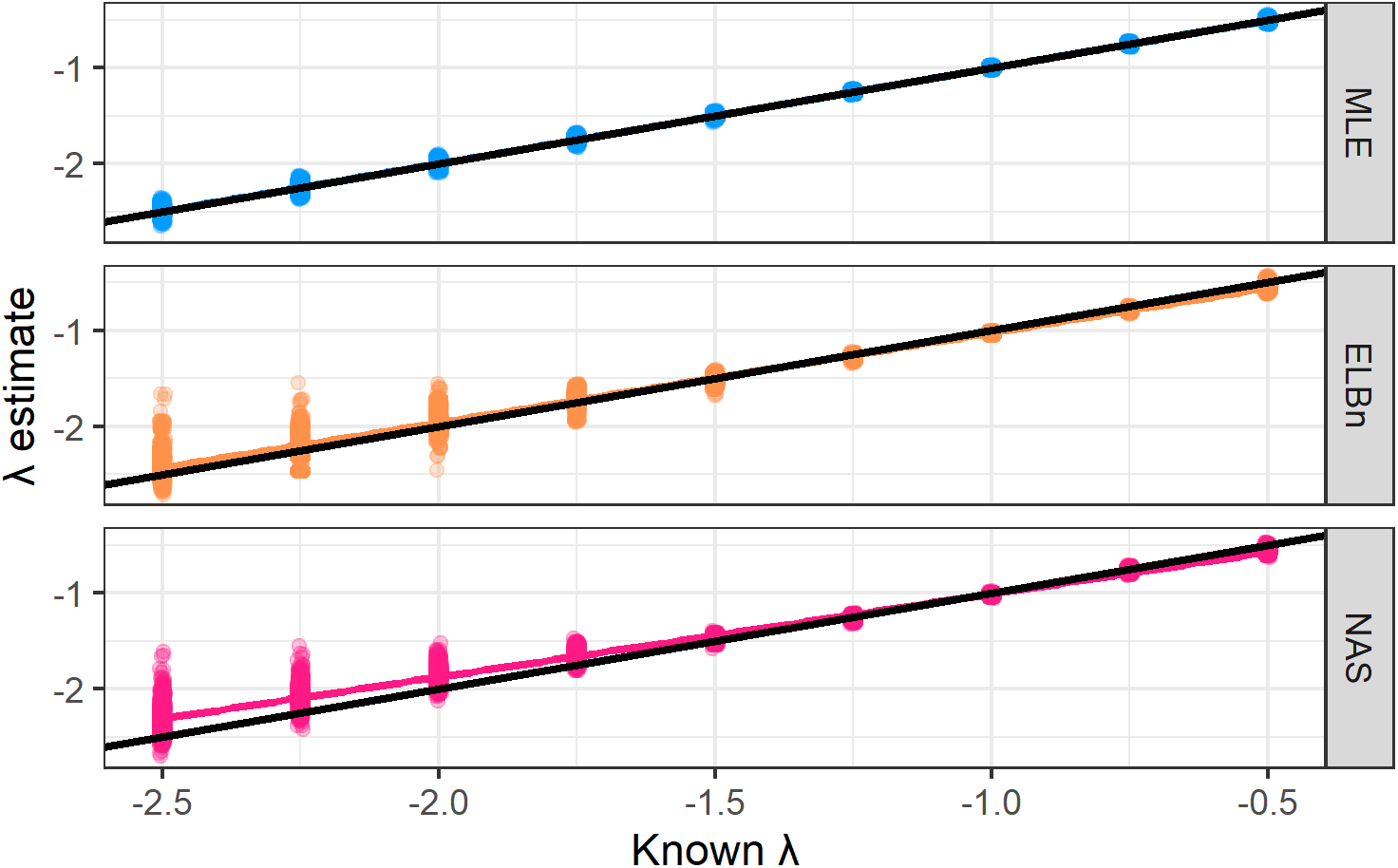
Figure 2. Plot of the known and estimated lambda from different methods (color). The dashed line is the 1:1 line, and almost perfectly matches the MLE (Blue) estimate. The two binning method cross the 1:1 line, meaning they systematically underestimate shallow lambdas, and over estimate steeper lambdas.

**Table 1** (*Submitted as additional file on biorxiv*.) Summary of three methods in recapturing the known values of lambda or the regression slopes simulated in this study. Performance is determined by comparinged by uncertainty (range of 95% CI’s) and the absolute distance of the model estimates from the known estimates (median and SD of the difference). Values are summarized across all n = 9000 or 6000 simulated data sets. See figures for more specific comparisons.

## Relationship across the hypothetical environmental gradient *λ* “windows”

Because of the different performance of the two binning methods at steep and shallow values of lambdas, we first investigated how well the methods capture a known relationship of *β*_*env*_ = - 0.5 across a hypothetical environmental gradient. The lambda “windows” are steep (−2.5, -1.5), medium (−2, -1) and shallow (−1.5, -0.5).

The MLE method (Figure 3, blue) recaptured the known slope value in each of the “windows.” with median absolute deviation of ∼0.01 units (Table 1). By contrast, the binning methods systematically overestimated the known slope (Figure 3), with median absolute deviations >4 times higher than the MLE. Similarly, uncertainty in the slope estimates derived from binning methods was twice that of the uncertainty in the MLE method (Table 1).

**Figure 3:**
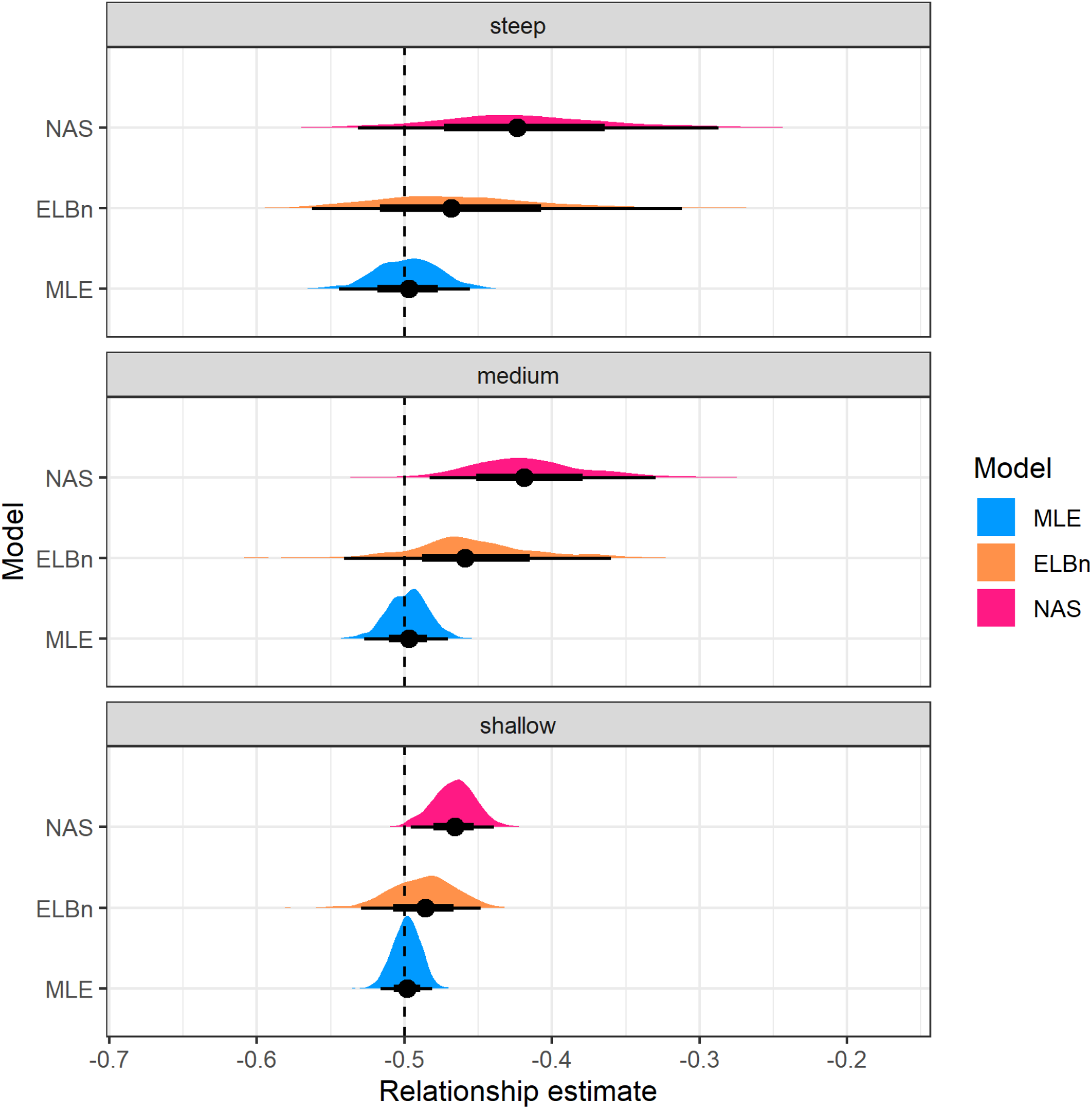
Figure 3. Distribution of relationship estimates (*beta*_*env*_) in three different “windows” of lambda values. The dashed vertical line is the known relationship value of -0.5.

### Varying the Known Relationship

All methods recaptured the correct sign of the slopes, yielding qualitative consistency (Figure 4). However, the binning methods again systematically underestimated the true value of the slope by ∼0.05 units (Figure 5). Likewise, uncertainty in the slope estimates was always greater in the binning methods, with the width of the distributions increasing with stronger relationships across a hypothetical gradient. By comparison, the MLE showed no evidence of bias and was always centered at the known value with relatively narrow variation.

**Figure 4:**
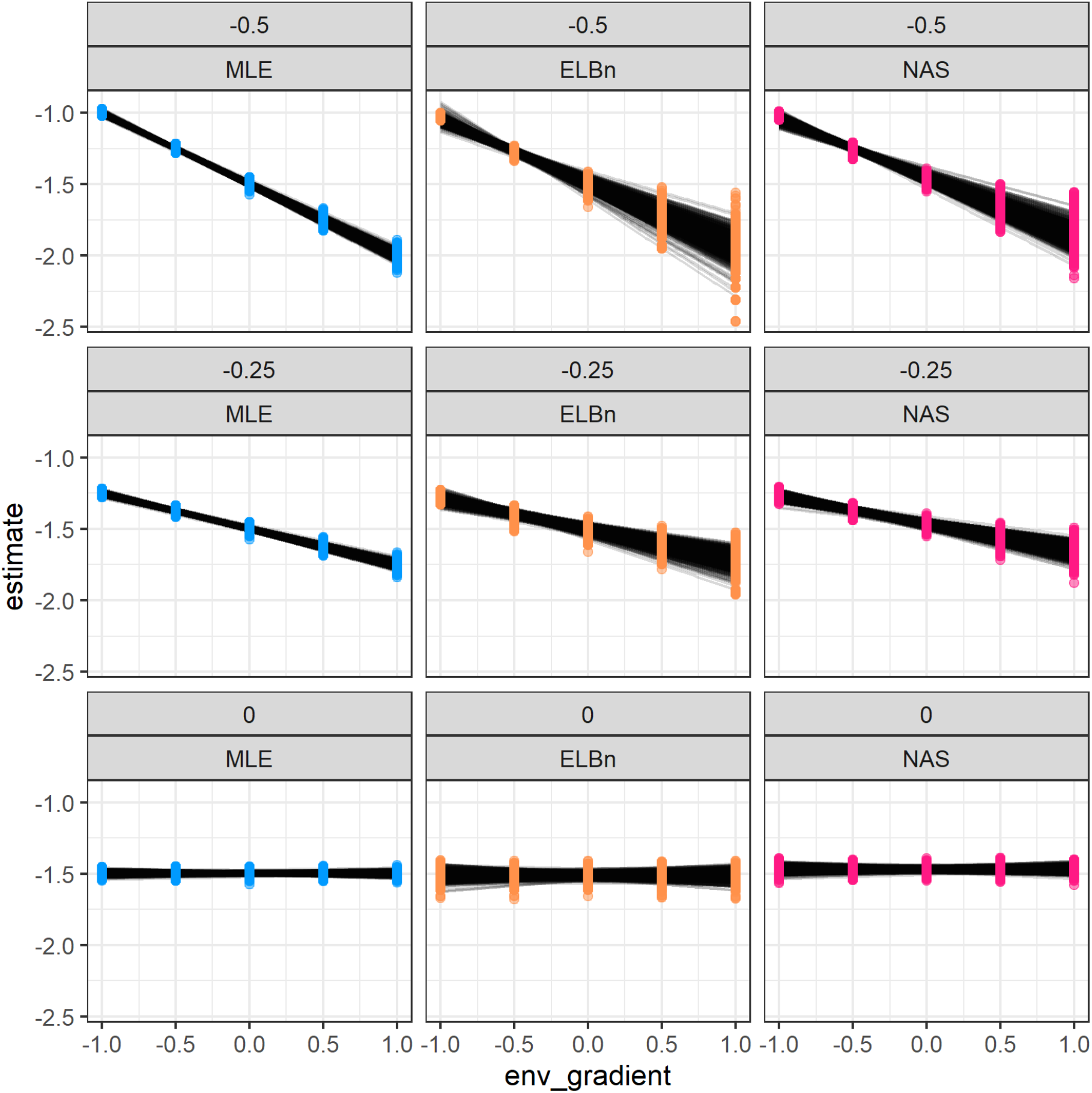
Figure 4. Individual regressions for each rep (N = 1000) and method (columns) for three different known relationship values (rows from bottom to top): 0, 0.25, and 0.5

**Figure 5:**
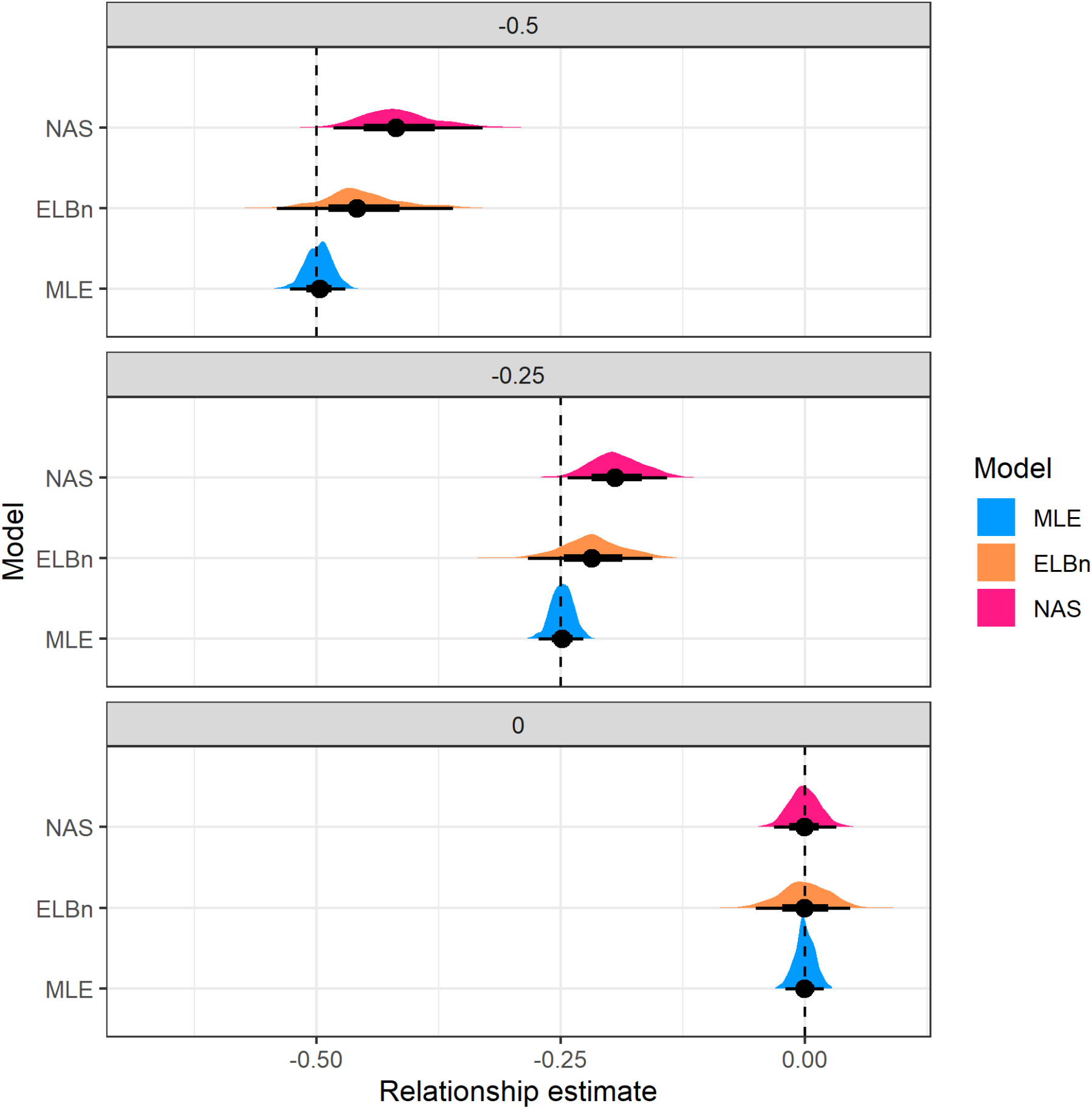
Figure 5. Distribution of relationship estimates (beta_1) when estimating from different known relationships

### Empirical data

Both empirical data sets yielded similar results to the simulated data; the direction of the coefficients (i.e., *β*_1_ coefficients) were the same among methods, with positive slopes across the AMD gradient (Fig. 6A) and negative slopes across the temperature gradient. However, the magnitude of change differed between methods. The AMD slopes ranged from ∼0.62 with MLE to ∼0.78 with NAS (Figure 6b), while temperature slopes ranged from -0.0058 with MLE to -0.0019 with ELBn (Figure 6d). As with simulated data, slope uncertainty (± 1 SD) was larger in the binning methods, particularly the ELBn method (Fig. 6B). Likewise, the size spectra parameters consistently increase (become steeper) with increasing temperature across the NEON sites (Fig. 6C).

**Figure 6:**
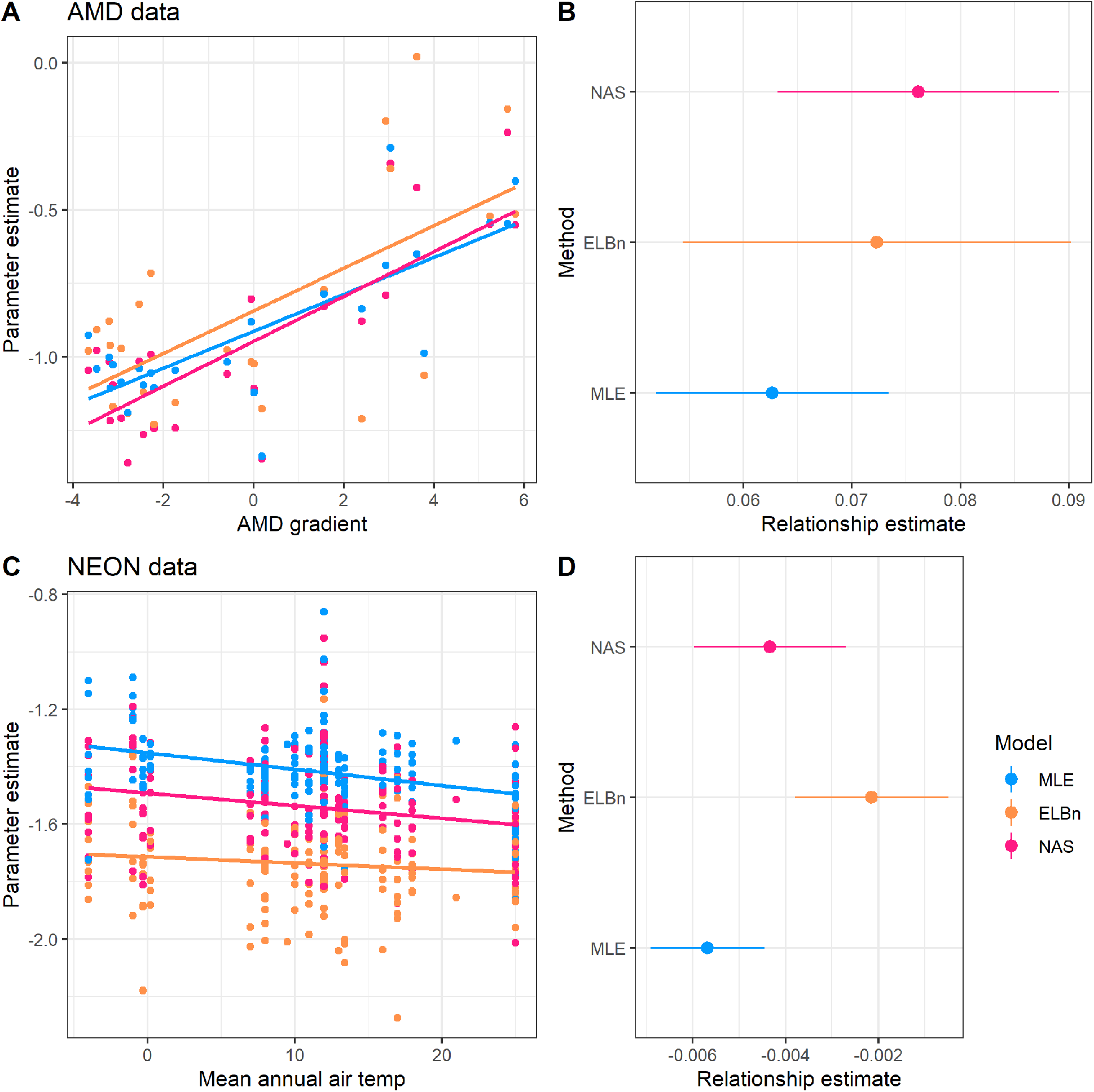
Figure 6. of empirical data estimates. All of the methods estimate the same sign of the relationship, but the estimates from the binning methods are generally greater than the MLE estimates.

## Discussion

The relationship between body size and abundance has been extensively studied in a wide range of taxa inhabiting both terrestrial and aquatic ecosystems (reviewed by Brown (1995), and White et al. (2007)). Empirical data shows generally consistent patterns and can be explained by the metabolic theory of ecology (Brown et al. 2004). Measuring parameters describing the decline in abundance with increasing body size in communities is being done with increasing frequency across ecology. Previous work has investigated the accuracy and inherent biases associated with different estimation methods (White et al. 2007, Edwards et al. 2017, Edwards et al. 2020). However, the extent to which these inaccuracies and biases compound across environmental gradients remains uncertain, making it difficult to detect variation in size spectra parameters across environmental gradients with confidence. The most important outcome of our work is that binning methods not only generate biased *λ* values for individual data sets, but that bias carries over to affect the parameters of subsequent regressions that use those *λ*’s as response variables. This makes it challenging to understand how *λ* varies in response to environmental gradients if binning is used to estimate size spectra exponents.

Binning methods are easy to use and interpret, which most likely accounts for their wide use in ecological studies. However, aggregating individuals into logarithmic bins removes a large amount of information within the data by collapsing body size variation into a single value within each bin. For example, all individuals placed into a bin that ranges from 2-4 grams of mass are all treated as having a mass of 3 grams, the midpoint of that bin. Likewise, a single abundance value is taken for each bin, despite that fact that there is almost certainly variation in the abundance of individuals that weigh ∼2, ∼3, or ∼4 grams. Moreover, the number of bins that can be produced by any dataset is limited most often to less than six. That means a subsequent log-log regression used to estimate *λ* would contains n = 6 data points, even though there likely 100’s or 1000’s of individual body sizes available in a typical body size data set. By homogenizing the data in this way, binning methods produce noisier results than MLE, since binning is akin to deleting information that the model could otherwise use. By contrast, the MLE uses all the individual body size data to directly estimate *λ*, meaning that it not only produces more accurate estimates, but does so with less uncertainty than binning, even when the underlying data sets are identical.

Although there were differences in the value of the empirical relationship parameters, they were generally in the correct direction and of a similar magnitude. This suggests that previously reported changes in size spectra parameters across environmental gradients and in experimental manipulations are plausible. However, the biases and inconsistencies in the estimates of both lambda and environmental response parameters presented here suggest that it may be difficult if not impossible to directly compare the relative changes across different published studies which use different methods.

The publication of individual body size data with future studies of size spectra would greatly aid in our ability to generalize changes to this fundamental aspect of community organization across spatiotemporal scales and in response to environmental conditions.

### Concluding Remarks

The MLE method outperformed both of the binning methods under nearly any measure. With the publication of the *sizeSpectra* package (Edwards et al. 2017), producing MLE estimates of size spectra parameters is a relatively easy task. Therefore, we recommend using it in all future studies of size spectra relationships rather than binning.

We reiterate the recommendations of White et al. (2007), Sprules and Barth (2015), and Edwards et al. (2017) to estimate size spectra relationships using MLE methods due to their superior performance in nearly every context. Furthermore, we strongly encourage authors to publish individual size data whenever possible. This will allow for the consistent re-analysis of existing data sets as methodologies develop and improve. This will aid in the ability for size spectra work to be synthesized between research groups and across scales.

## Supplementary material

### *λ* and relationship estimates

In the main analysis, we simulate body size data from bounded power law distributions while varying the *λ* exponent which describes the distribution. For the results presented in the main text, we held the number of sites at 5, scaled the environmental gradient from -1 to 1, and set the minimum and maximum body sizes to 0.0026 and 1.2 ∗ 10^3^ respectively. Here, we plot the results of varying the number of sites (3, 10), increasing the scale of the environmental gradient (−100 to 100) and decreasing the range of body sizes (min = 1, max = 100). Generally, the results reported in the main manuscript are robust to changing these parameters: the MLE estimate is nearly always closer to the known parameters, and the variation in these estimates is usually smaller than the binning methods.

### Varying number of sites

#### 10 sites

**Figure S1:**
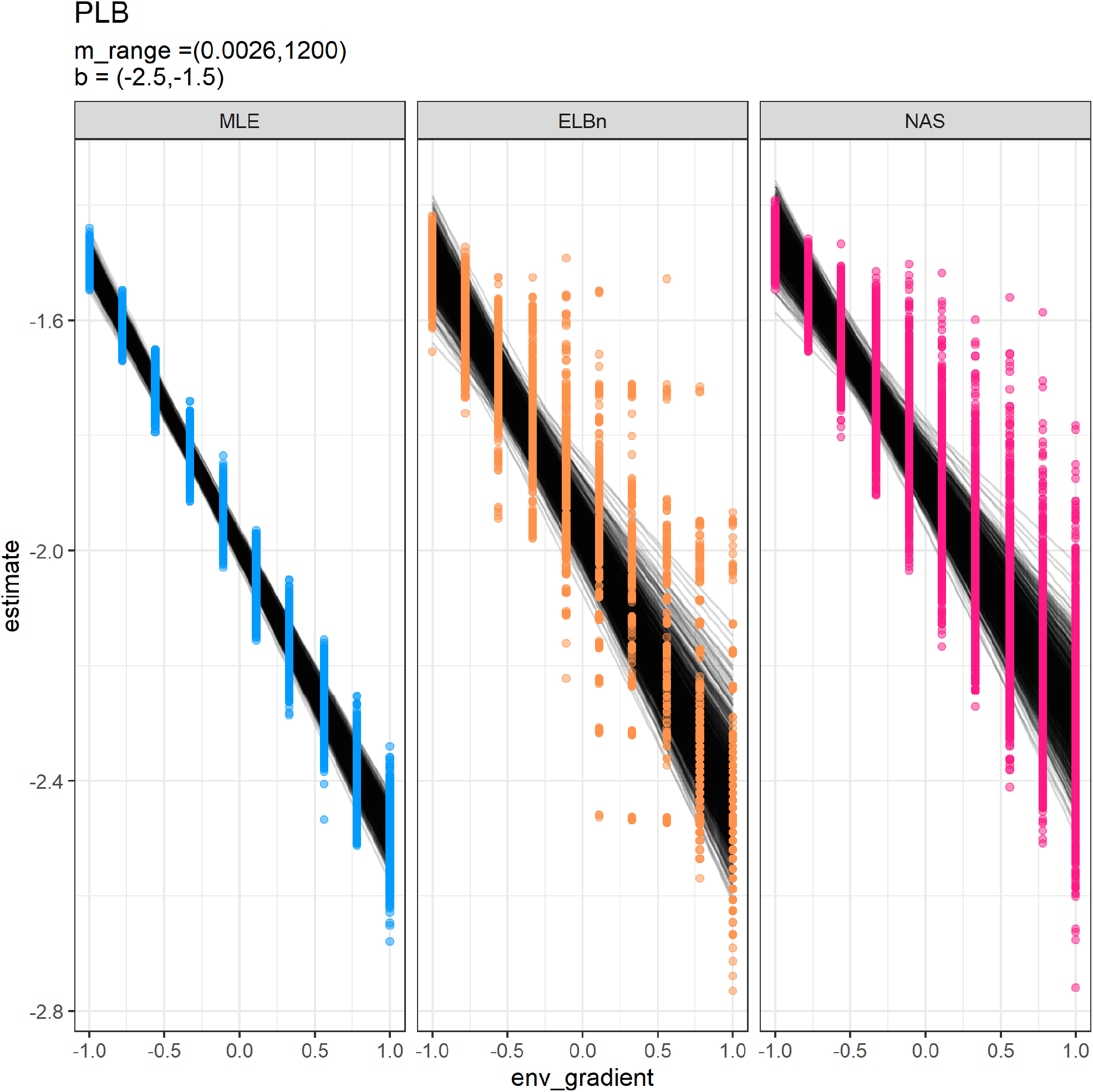
Individual regressions for ten sites across a hypothetical gradient with a known relationship of 0.5. All other parameters are the same as in the main analysis

**Figure S2:**
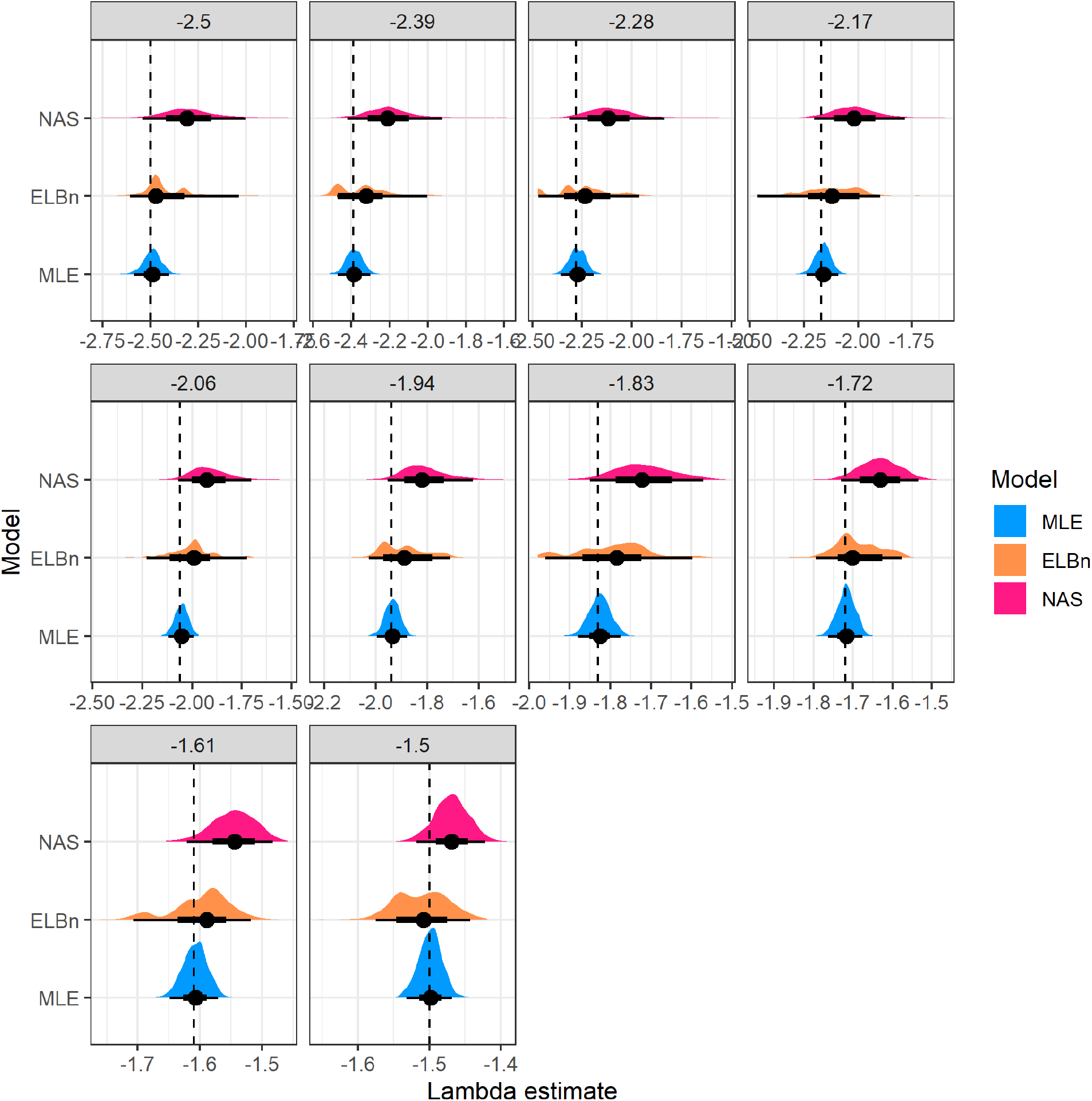
Distribution of estimated *λ* coefficient for ten sites across a hypothetical gradient with known values. All other parameters are the same as in the main analysis

**Figure S3:**
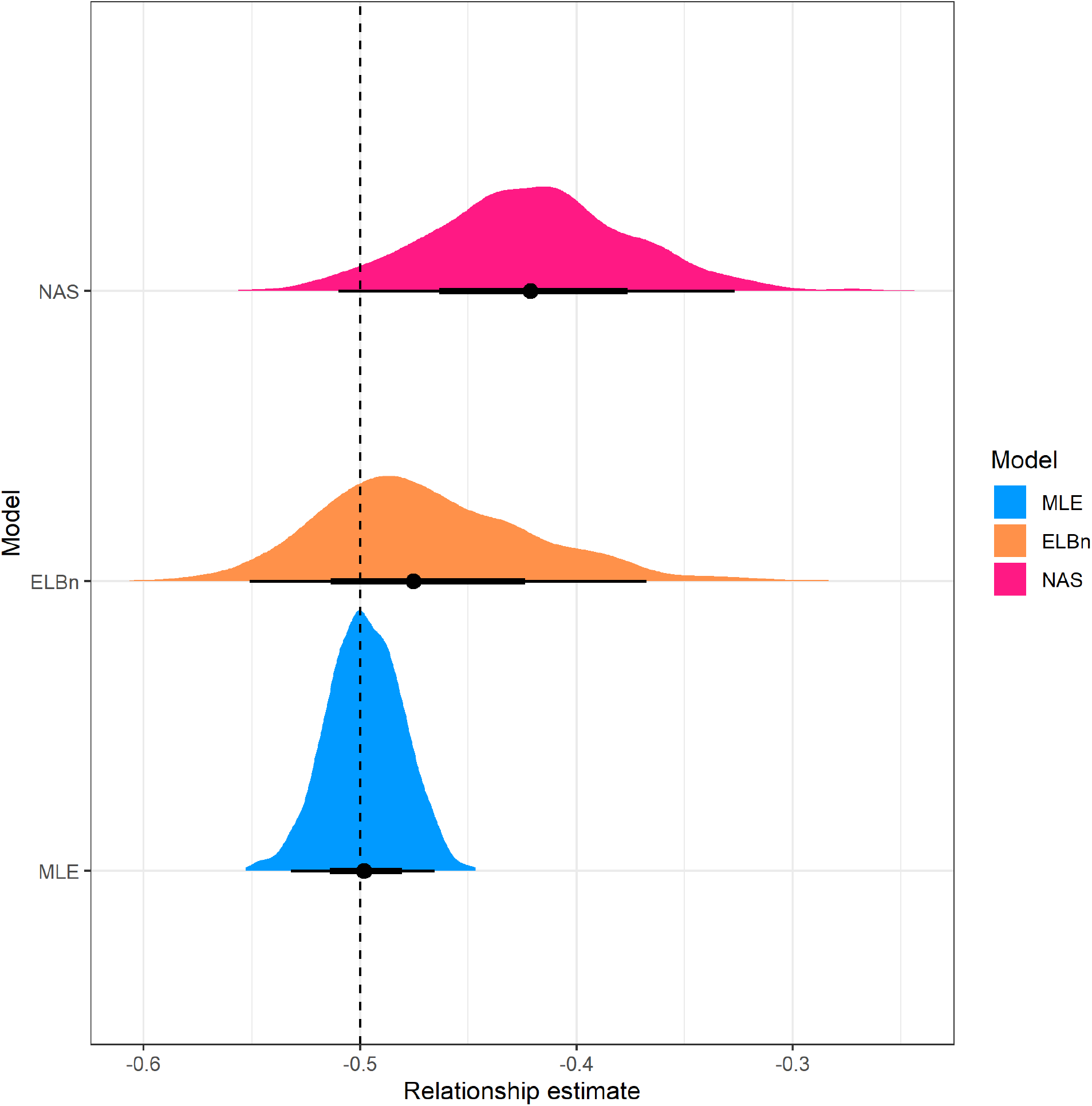
Distribution of estimated relationship (*β*_1_) coefficient’s for ten sites across a hypothetical gradient with known value of 0.5. All other parameters are the same as in the main analysis

#### Three sites

**Figure S4:**
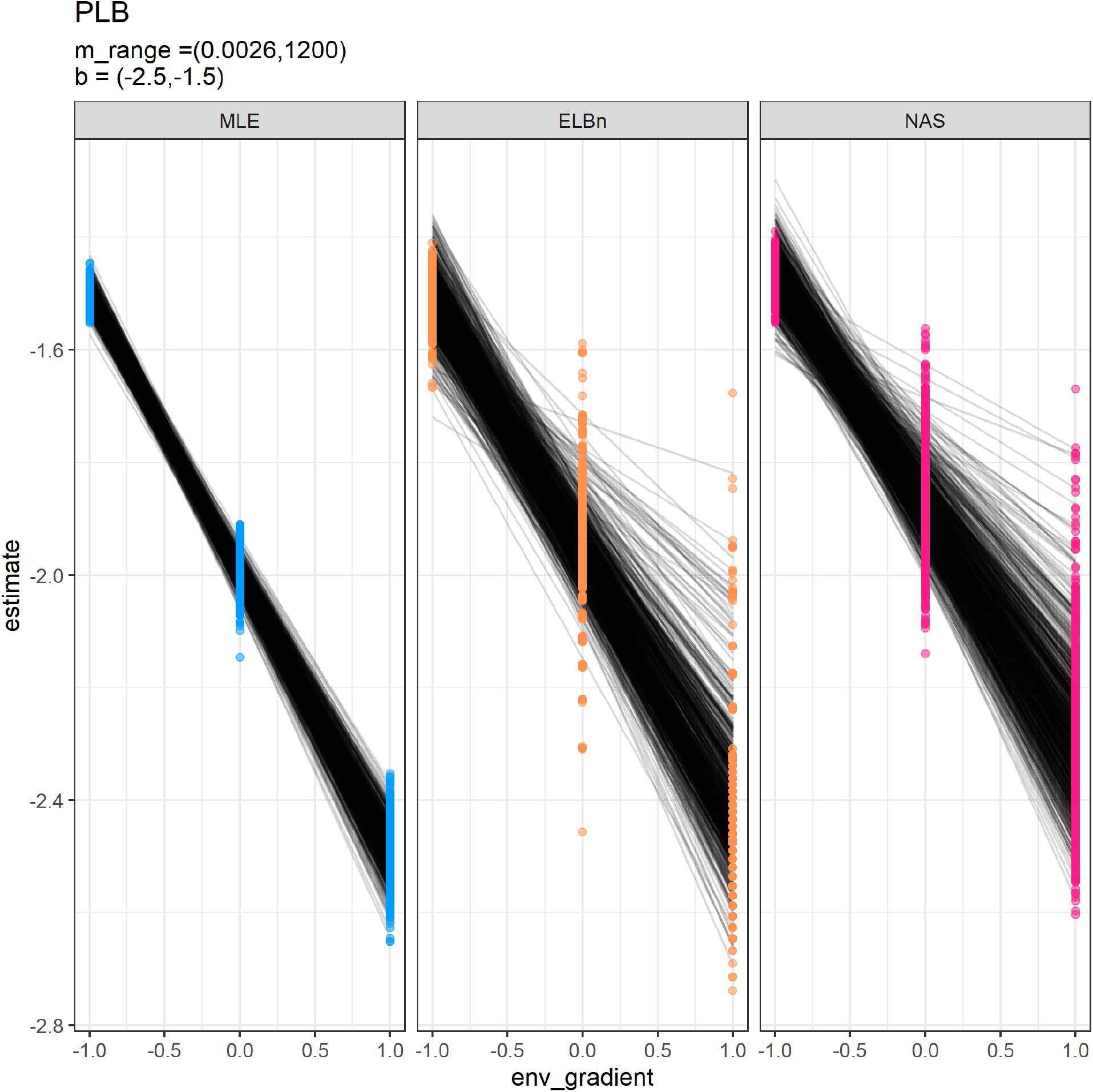
Individual regressions for three sites across a hypothetical gradient with a known relationship of 0.5. All other parameters are the same as in the main analysis.

**Figure S5:**
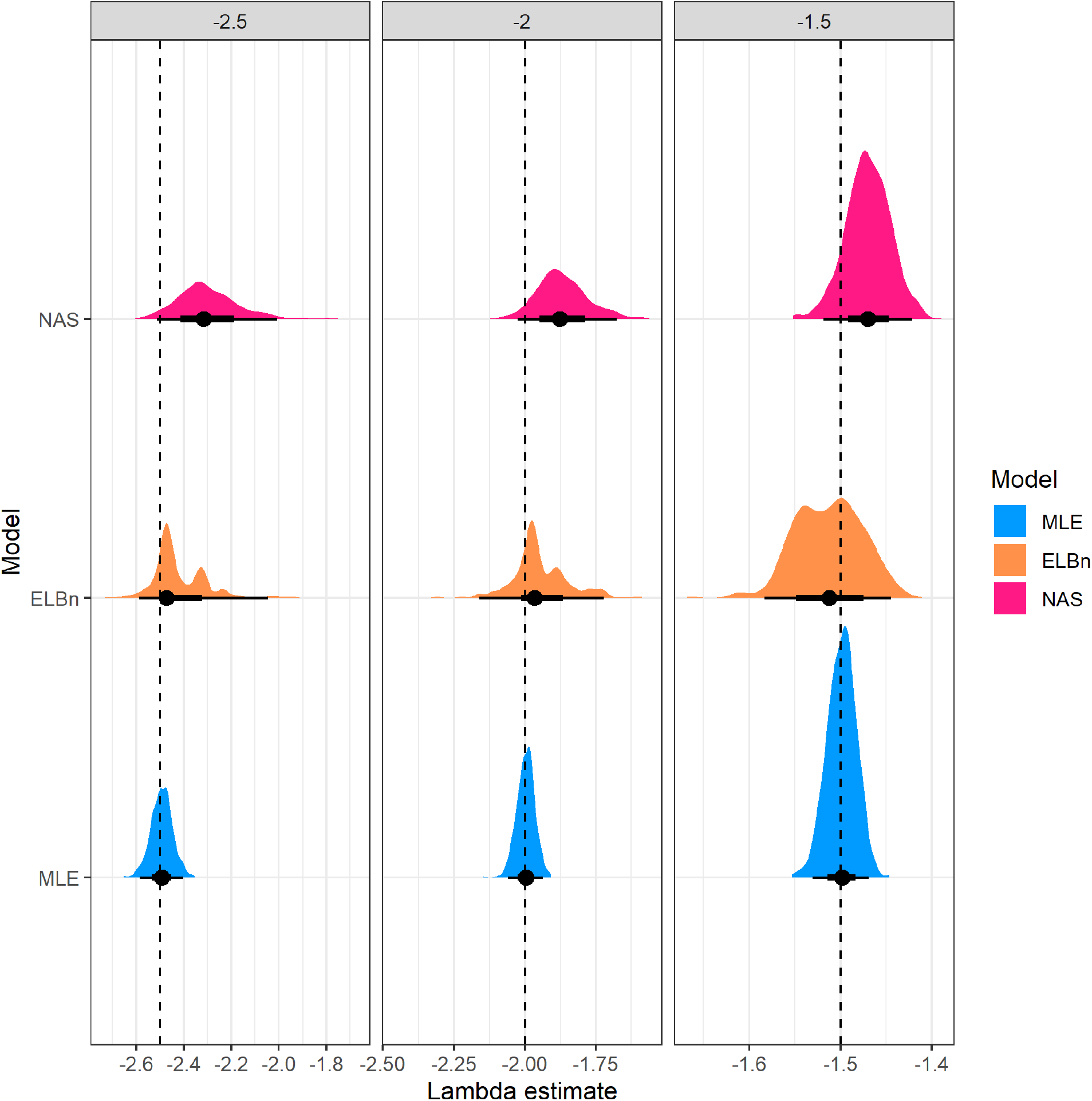
Distribution of estimated *λ* coefficient for three sites across a hypothetical gradient with known values. All other parameters are the same as in the main analysis.

**Figure S6:**
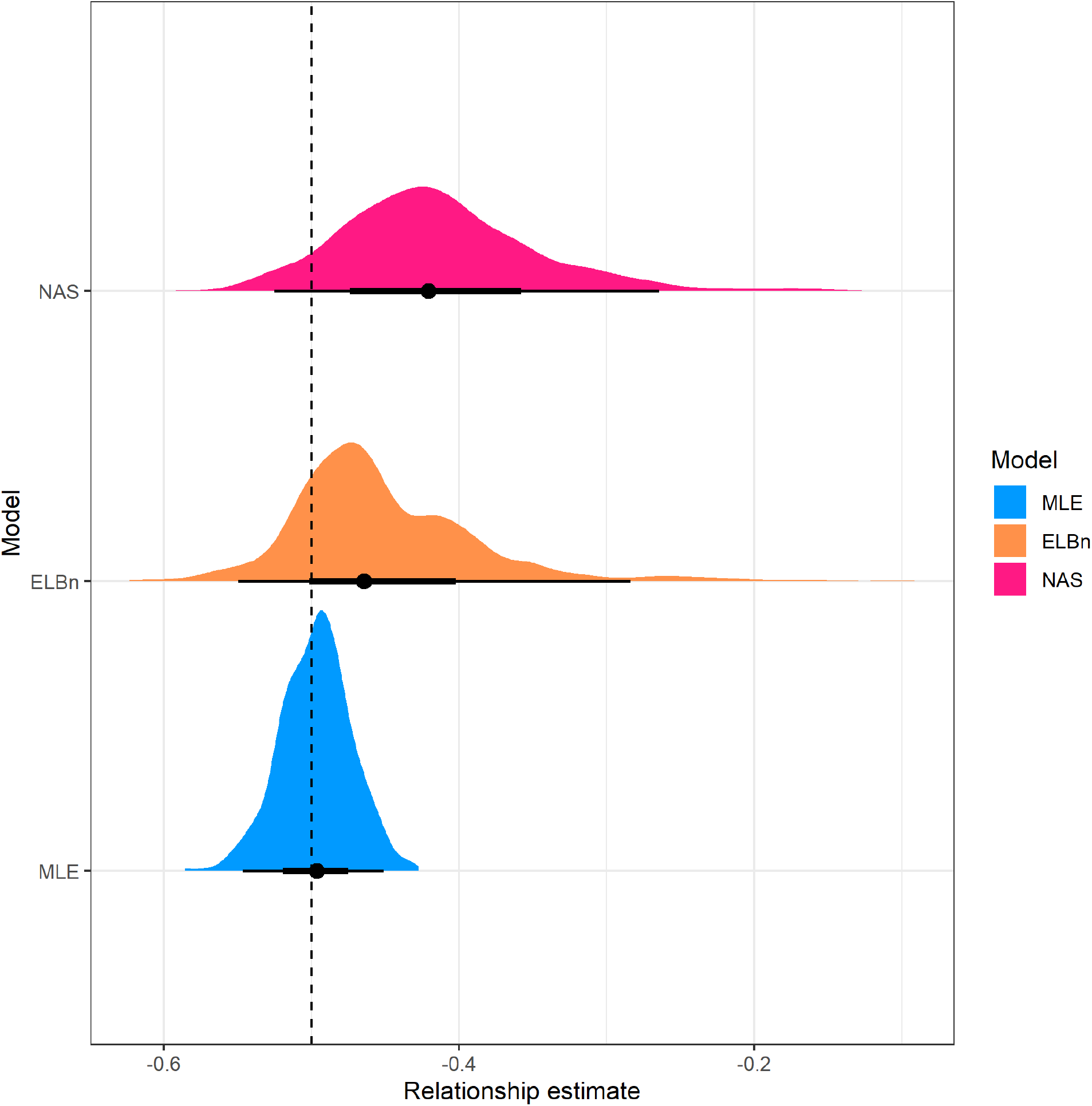
Distribution of estimated relationship (*β*_1_) coefficient’s for three sites across a hypothetical gradient with known value of 0.5. All other parameters are the same as in the main analysis

### Large environmental gradient

**Figure S7:**
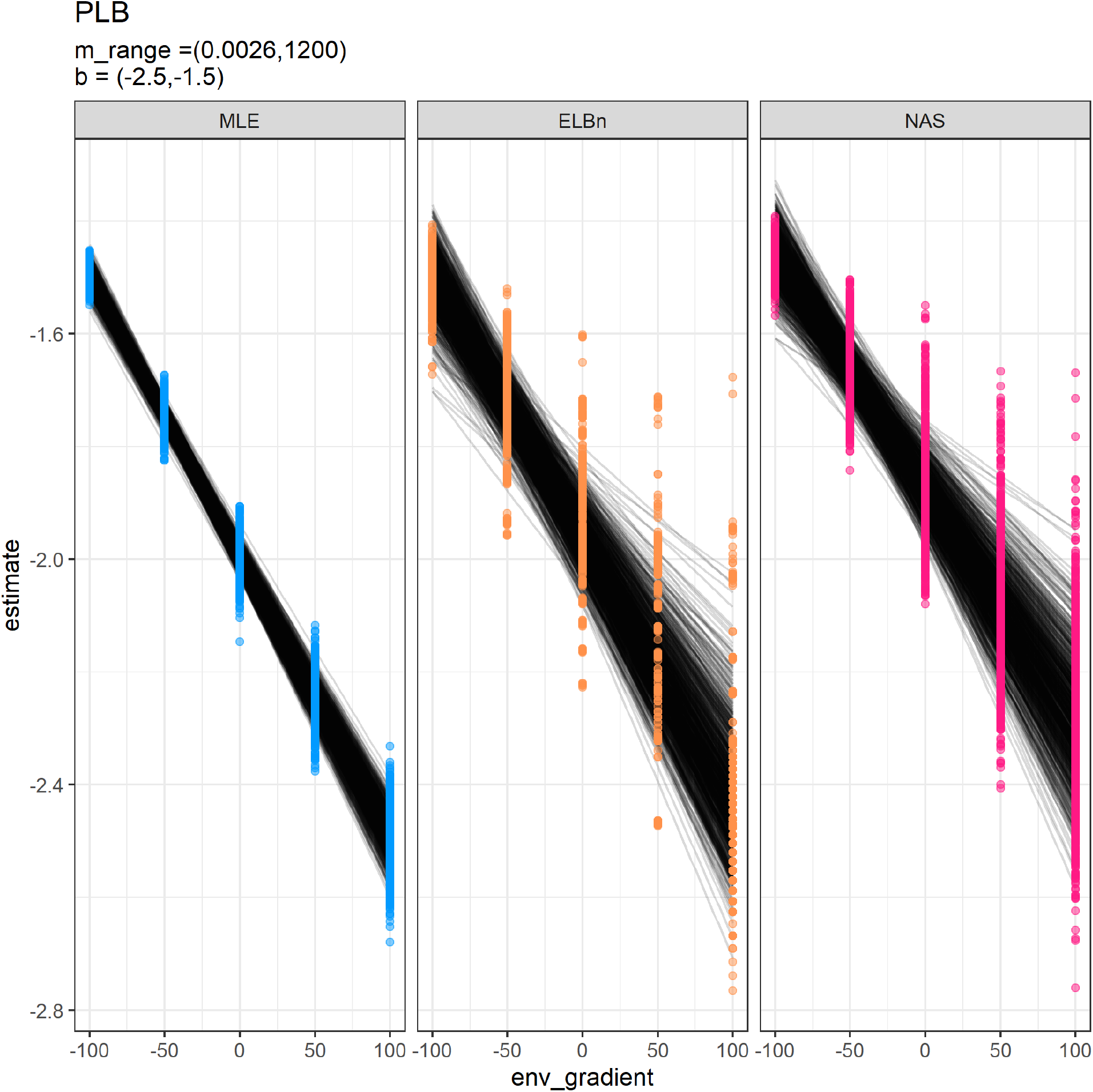
Individual regressions for five sites across a hypothetical gradient with a known relationship of 0.5. Range of environmental values (*x*-axis) increased to be -1000, to 1000.

**Figure S8:**
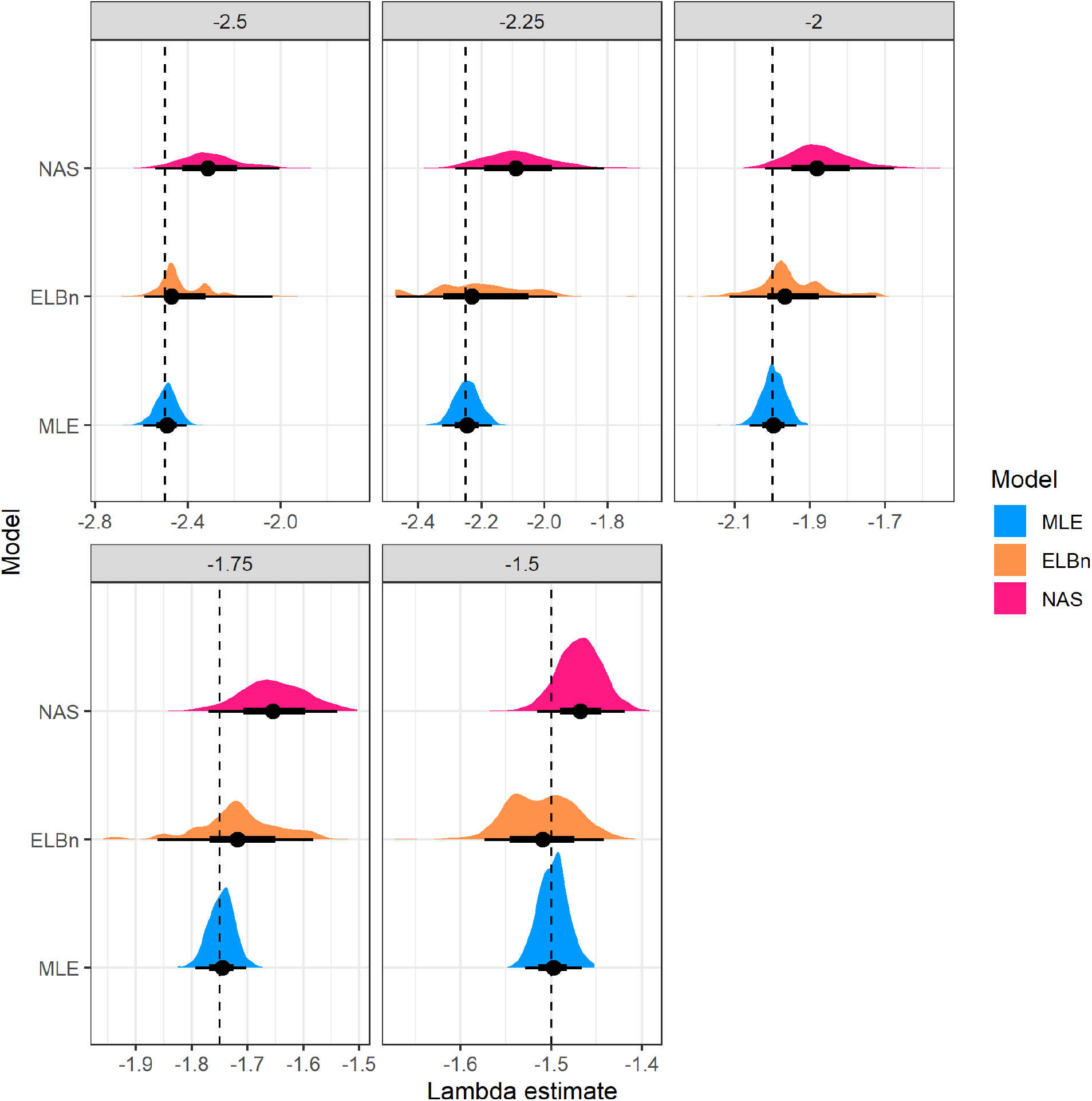
Distribution of estimated *λ* coefficient for five sites across a hypothetical gradient with known values. Range of environmental values (*x*-axis) increased to be -1000, to 1000.

**Figure S9:**
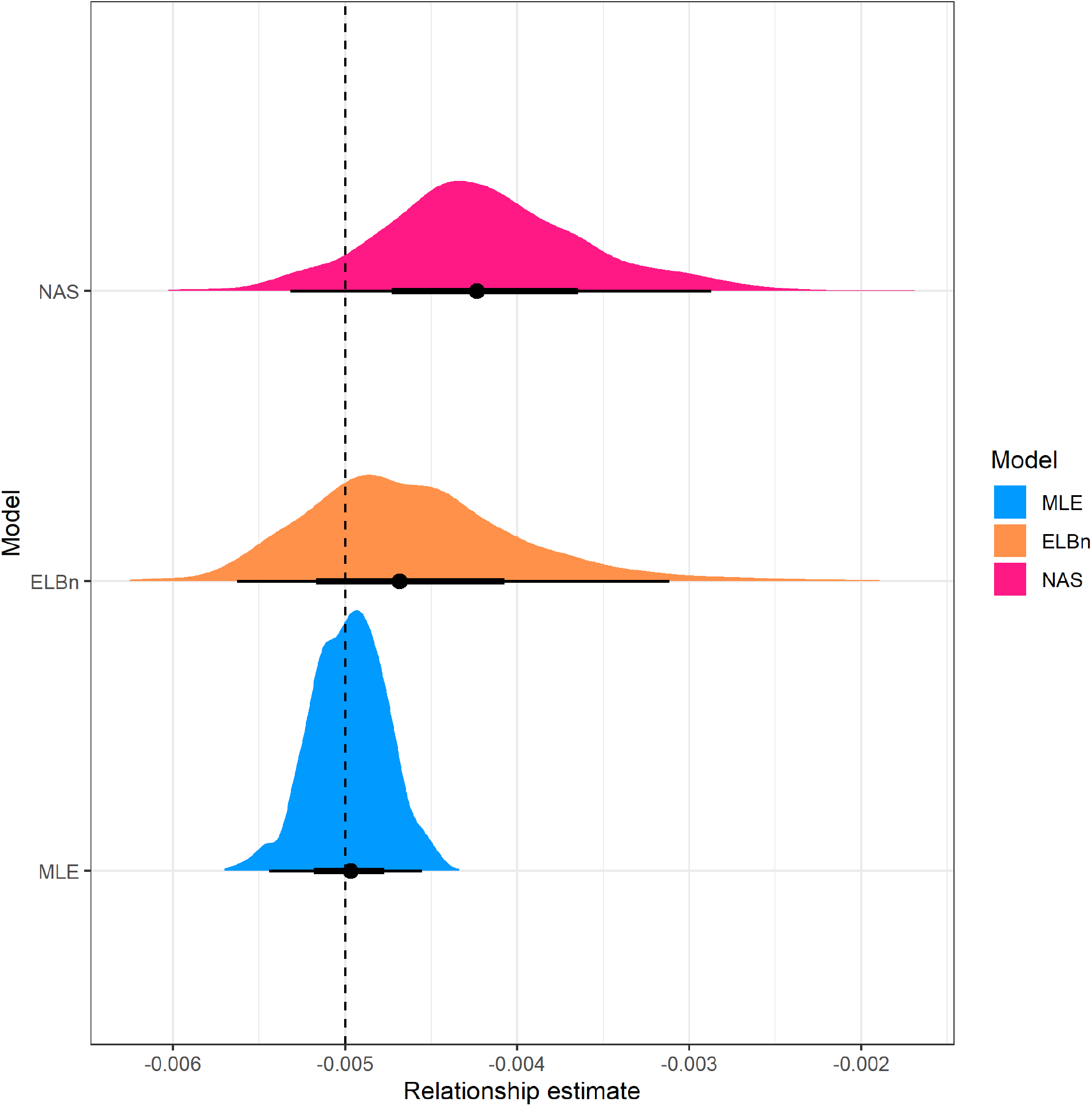
Distribution of estimated relationship (*β*_1_) coefficient’s for five sites across a hypothetical gradient with known value of 0.5. Range of environmental values (*x*-axis) increased to be -1000, to 1000.

### Range of body sizes, *M*

**Figure S10:**
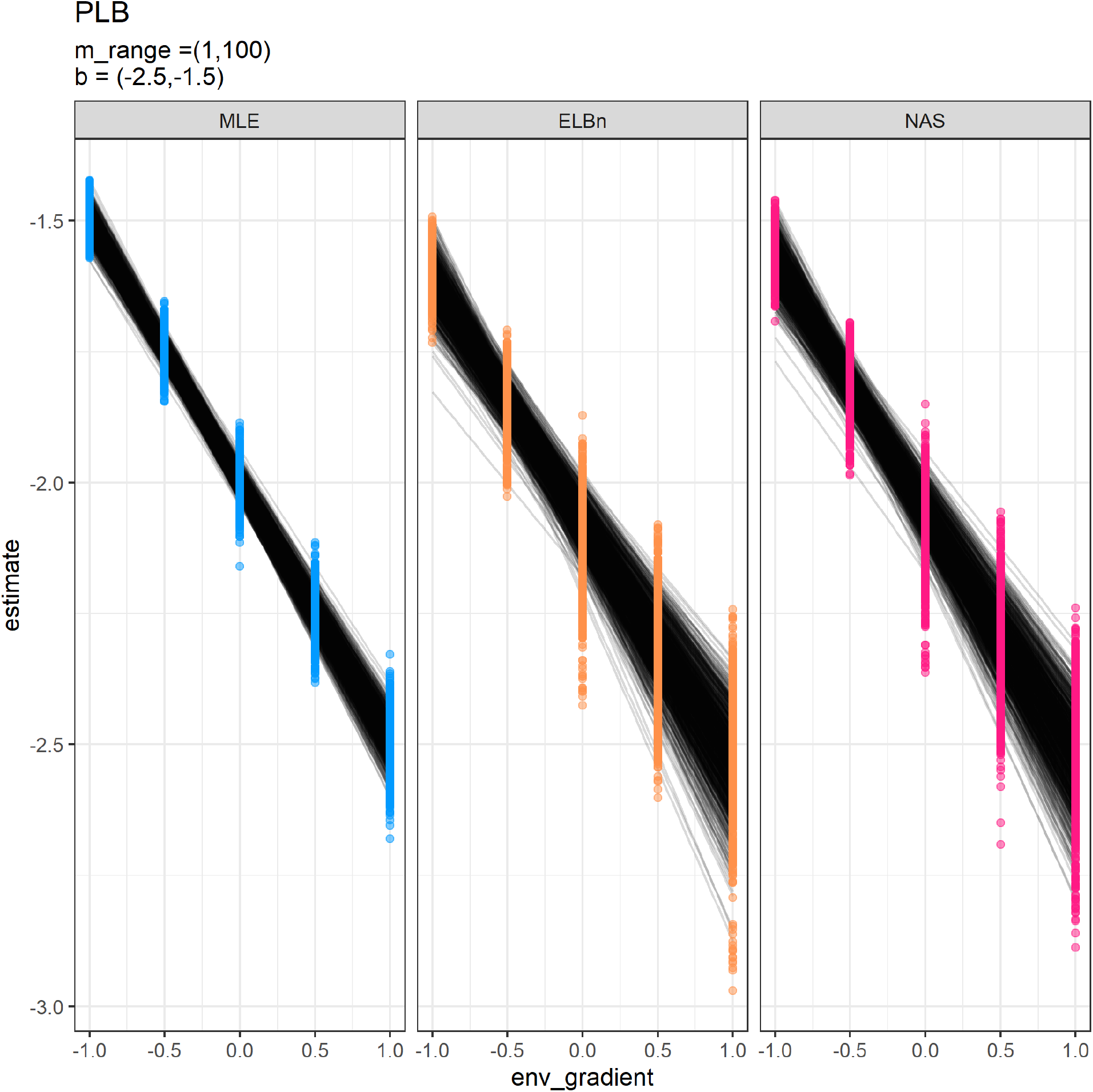
Individual regressions for five sites across a hypothetical gradient with a known relationship of 0.5. Range of body sizes is reduced and is from 1, to 100.

**Figure S11:**
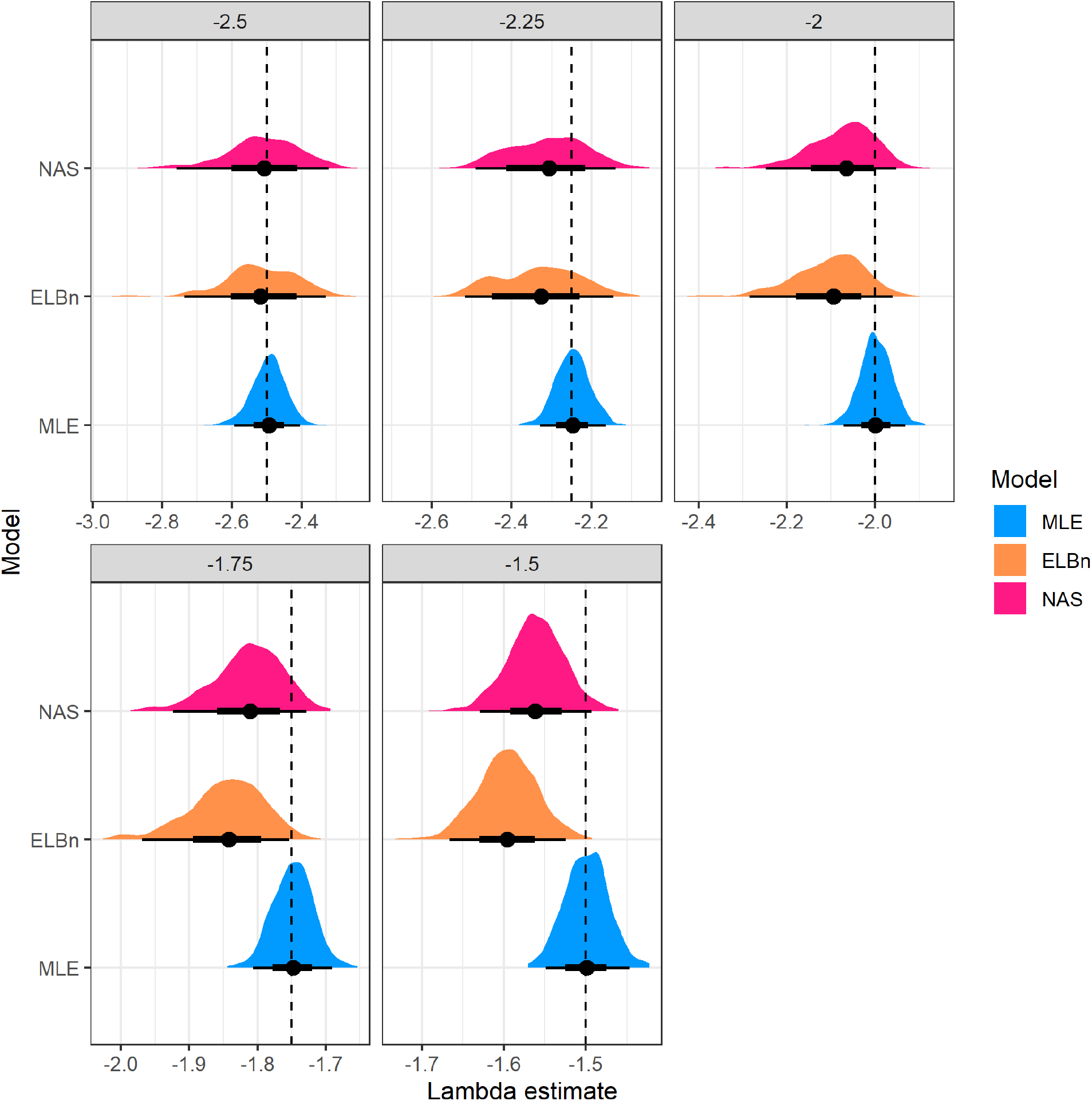
Distribution of estimated $\lambda$ coefficient for five sites across a hypothetical gradient with known values(dashed line). Range of body sizes is smaller than main anaysis and ranges from 1, to 100.

**Figure S12:**
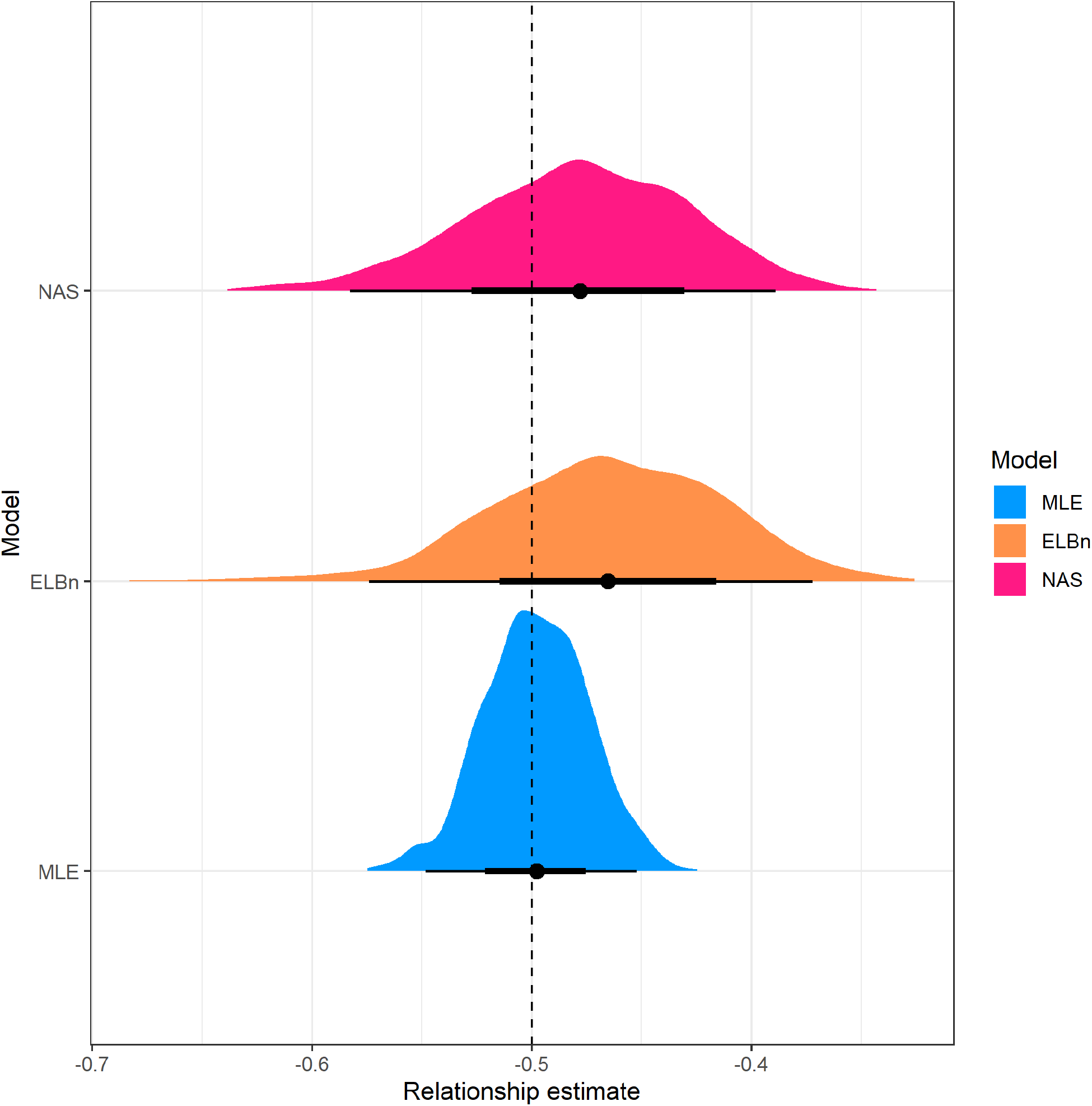
Distribution of estimated relationship (*β*_1_) coefficient’s for five sites across a hypothetical gradient with known value of 0.5. Range of body sizes is reduced and is from 1, to 100.

### Sample size, *n*

The number of observations in our simulations may bias the results. Therefore, we repeated the simulations described above, but varied the sample size *n*. We tested values of *n* = 200, 500, 1000, 5000, 10000. Results of this analysis are presented in the Supplemental Information.

**Figure S13:**
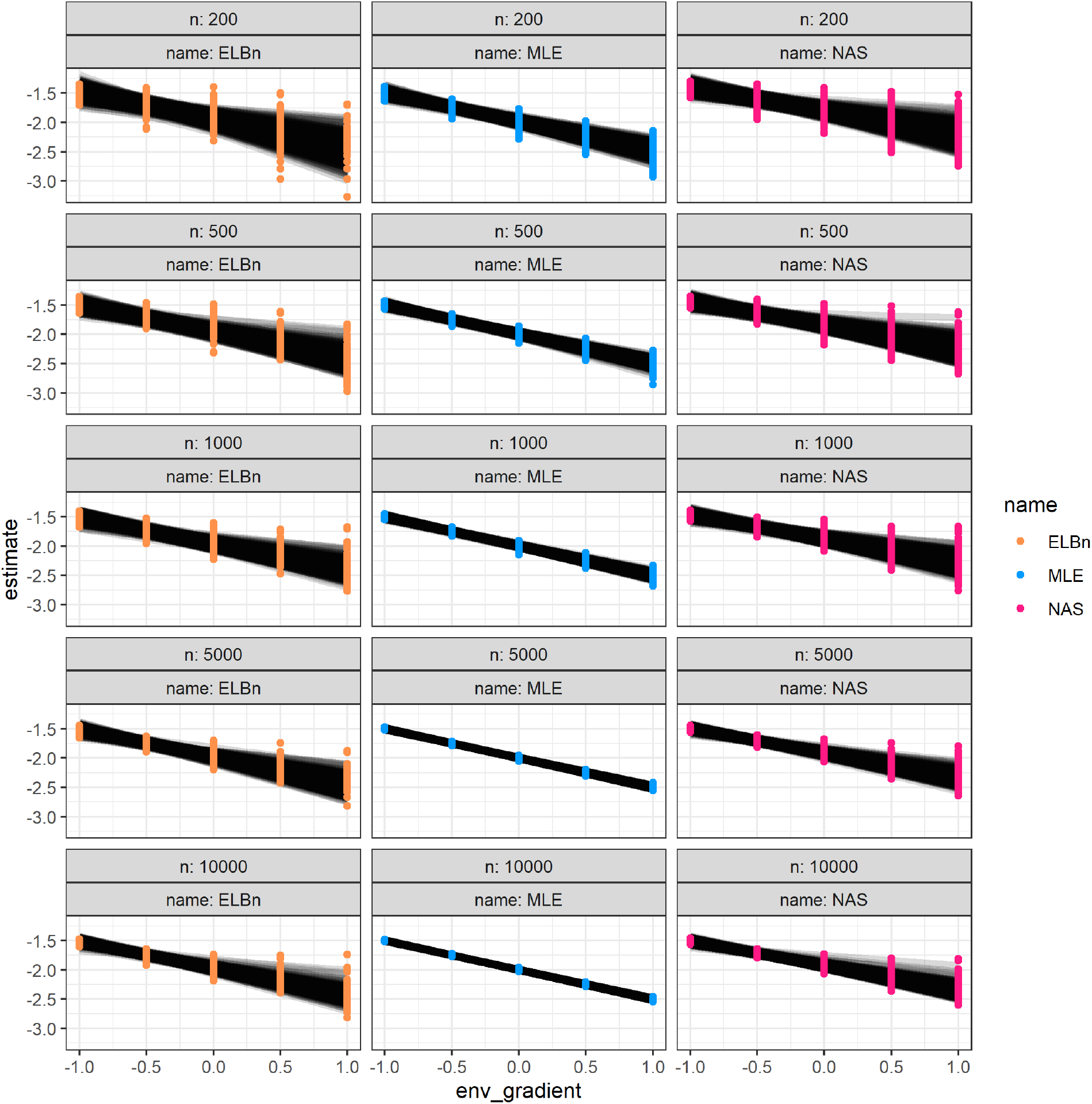
Individual regression estimates across the hypothetical gradient based on sample size (rows) and methodology used (columns). (match this figure to “new” style if we like that better)

**Figure S14:**
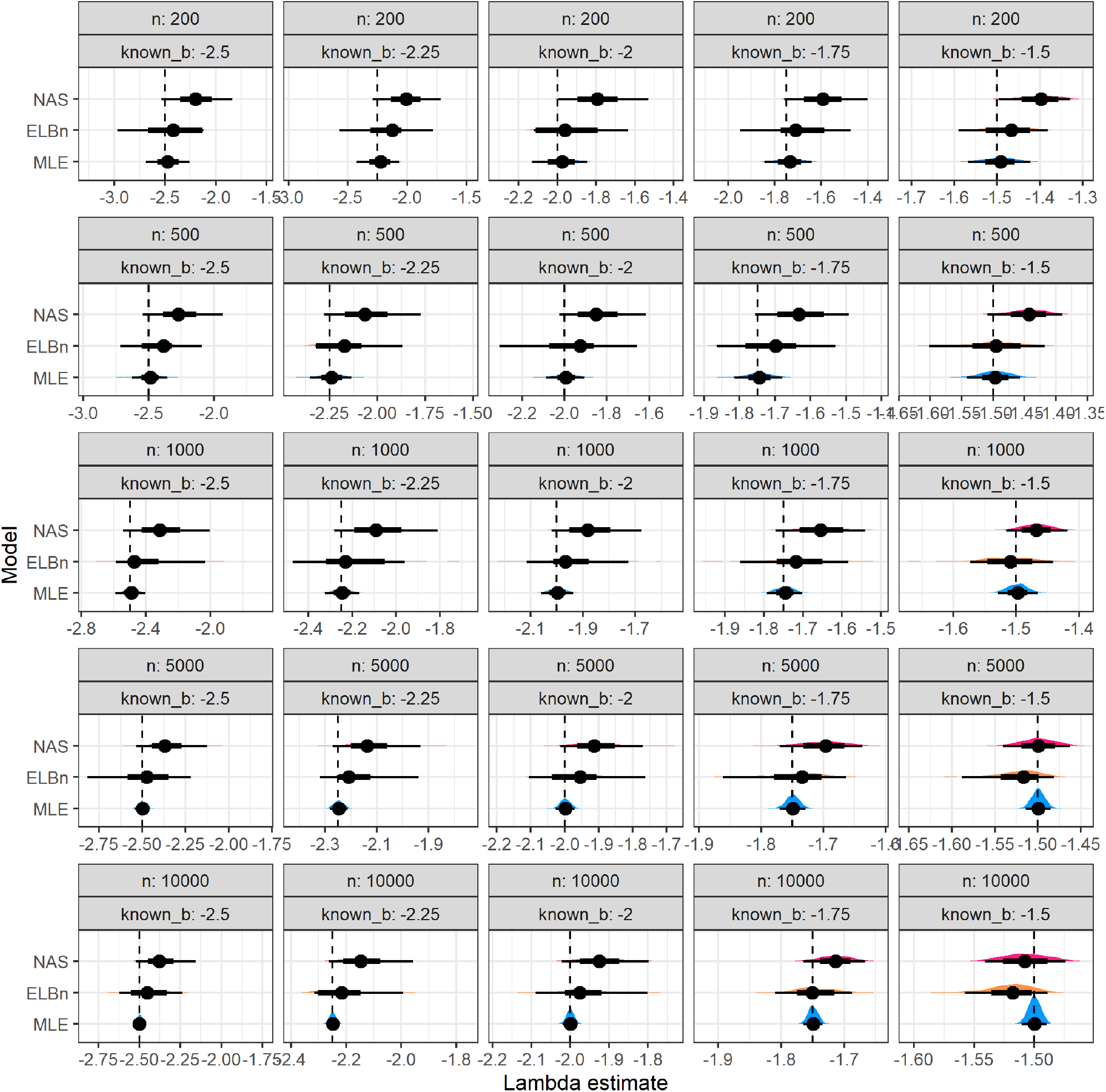
Distribution of size spectra parameter estimates. Vertical line is the known parameter (dashed line) wich describes the bounded power law distribution from which the body size estimates were sampled. As n increases (top to bottom) and *λ* increases (left to right), the accuracy of the estimate improves across all methods.

**Figure S15:**
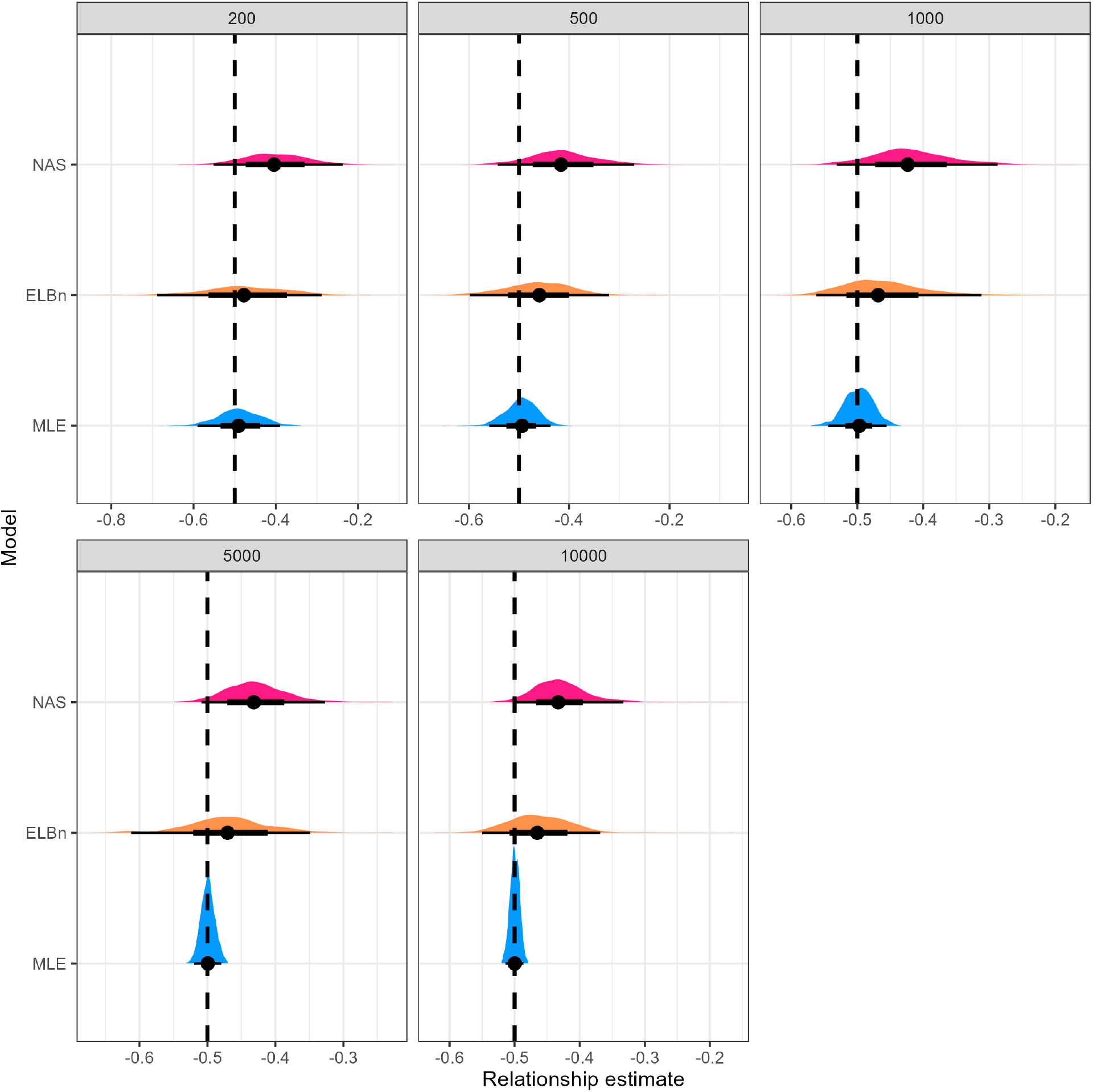
Distribtuion of the relationship coefficients with varying sample size

## Comparison with other published estimates

SI Table. This table shows published estimates of the variation in size spectra slopes (or exponents) in empirical studies. It is unclear how to directly compare estimates of the slope with different methods. However, the published estimates here range from ∼0.1 to 0.2 across the gradients studied. For comparison, the 2.5-95% quantiles around the relationship estimate for the MLE method were ∼0.1, whereas for the ELBn and NAS method they were ∼0.25 and ∼0.2, respectively. b_diff is the change in estimate (b-low-b-min). System refers to stream communities or mesocosm experiments. Method: MLE = maximum likelihod estimate, ELB = equal logarthmic binning, the number before indicates the number of bins used. The normalization process shifts the estimates by an absolute value of 1.0. Hence, direct comparison the relative change in normalized and non-normalized studies should not introduce any bias. The O’Gorman et al. 2017 study used average species size and abundance (Local Size Density Relationship, *sensu* White et al. 2007) as opposed to individual size distribution. These methods are related, but it is unclear how to directly compare estimates from each method.

Supplementary table comparing with other published results will be submitted as a separate file for formatting.

## Notes

https://datadryad.org/stash/dataset/doi:10.5061/dryad.v6g985s

https://data.neonscience.org/data-products/DP1.20120.001

https://github.com/Jpomz/detecting-spectra-differences

### Competing Interest Statement

The authors have declared no competing interest.

